# The Red Queen’s Crown: an evolutionary arms race between coronaviruses and mammalian species reflected in positive selection of the ACE2 receptor among many species

**DOI:** 10.1101/2020.05.14.096131

**Authors:** Mehrdad Hajibabaei, Gregory A. C. Singer

## Abstract

The world is going through a global viral pandemic with devastating effects on human life and socioeconomic activities. This pandemic is the result of a zoonotic coronavirus, Severe Acute Respirsatory Syndrom Coronavirus 2 (SARS-CoV-2) which is believed to have originated in bats and transferred to humans possibly through an intermediate host species (Zhou et al. 2020; Coronaviridae Study Group of the International Committee on Taxonomy of Viruses 2020). The virus attacks host cells by attaching to a cell membrane surface protein receptor called ACE2 (Ge et al. 2013; Zhou et al. 2020). Given the critical role of ACE2 as a binding receptor for a number of coronaviruses, we studied the molecular evolution of ACE2 in a diverse range of mammalian species. Using ACE2 as the target protein, we wanted to specifically test the Red Queen hypothesis (Dawkins and Krebs 1979) where the parasite and host engage in an evolutionary arms race which can result in positive selection of their traits associated to their fitness and survival. Our results clearly show a phylogenetically broad evolutionary response, in the form of positive selection detected at the codon-level in ACE2. We see positive selection occurring at deep branches as well as 13 incidents at the species level. We found the strongest level of positive selection in Tasmanian devil (*Sarcophilus harrisii*), donkey (*Equus asinus*), large flying fox (*Pteropus vampyrus*), Weddell seal (*Leptonychotes weddellii*), and dog (*Canis lupus familiaris*). At the codon-level, we found up to 10% of ACE2 codons are impacted by positive selection in the mammalian lineages studied. This phylogenetically broad evolutionary arms race can contribute to the emergence of new strains of coronaviruses in different mammalian lineages with a potential to transfer between species given the common binding receptor ACE2. Our study provides a molecular evolutionary perspective to the current pandemic and sheds light on its evolutionary mechanisms.

*“Nothing in biology makes sense except in the light of evolution” (Theodosius Dobzhansky)*

## Introduction

Since Dec. 2019 a new strain of coronavirus, SARS-CoV-2, has generated a global pandemic with millions of individuals infected and approximately 300K dead (as of May 14^th^, 2020; World Health Organization, n.d.). With no cure or vaccine, this virus has had a drastic impact on human societies all over the world, especially as many countries have imposed rigid quarantine-like restrictions to limit human-human interactions and transmission of the virus. This virus causes coronavirus disease-19 (COVID-19) (World Health Organization, n.d.). In a majority of cases the infection manifests itself in a mild manner, from asymptomatic to mild upper respiratory tract symptoms. In a small proportion of individuals, especially older patients and those with comorbidities, it can lead to severe pneumonia and various complications such as lung injury, organ failures and eventually death (W. Guan et al. 2020). The main mode of transmission is through direct human contact via infectious body fluids but there is evidence of indirect transmission through contaminated surfaces and objects as well as airborne droplets or aerosols (World Health Organization, n.d.).

SARS-CoV-2 belongs to family Coronaviridae. These are ancient RNA viruses, with crown-like spike proteins on their capsid (hence the name *corona*, “crown” in Latin) many of which can generate different types of infections in different animal species—including human (Cui, Li, and Shi 2019). If there is close interaction between different species, coronaviruses can jump from one species to the other. They can spread to new hosts including companion animals, poultry, livestock, wild animals, and humans (Cui, Li, and Shi 2019). The first SARS-CoV originated in the Guangdong Province of China in 2002. Research showed that this virus originated in bats and was transferred to humans through an intermediate host species, possibly a palm civet (Y. Guan 2003). SARS-CoV caused over 800 deaths mostly in China, Hongkong, and Canada. Ten years later, in 2012, another CoV was detected in Saudi Arabia. This virus was called Middle East Respiratory Syndrome (MERS) CoV, which is believed to have originated from bats and transferred to humans through camels (Zaki et al. 2012). And there is evidence that the current pandemic coronavirus, SARS-CoV-2 has a very similar origin: first detected in Wuhan, China and either directly or through an intermediate host (e.g., a pangolin species) it has “jumped” from bats to humans and has gained human to human transmission (Andersen et al. 2020; T. Zhang, Wu, and Zhang 2020; Zhou et al. 2020).

Some coronaviruses including SARS-CoV and SARS-CoV-2 penetrate cells through binding to a receptor called Angiotensin Converting Enzyme 2 (ACE2) (Wenhui Li et al. 2003; Zhou et al. 2020). ACE2 is attached to the cell membranes in the lungs, arteries, heart, kidney, and intestines and is primarily involved in lowering blood pressure by catalysing the hydrolysis of angiotensin II into angiotensin. It has been shown that the SARS-CoV and SARS-CoV-2 receptor binding domain, spike S1 protein, directly binds to ACE2 and that this binding affinity is higher in SARS-CoV-2 compared to SARS-CoV (Wan et al. 2020). It has been suggested that ACE2 is both the critical entry receptor for SARS-CoV-2 and that it is also important in protecting the lungs from injury (H. Zhang et al. 2020). Highlighting the importance of the receptor, a new study has suggested a synthetic ACE2 can potentially be used as a therapy by binding to viral particles and preventing them from attacking endogenous ACE2 in cells (Monteil et al. 2020).

Understanding the evolutionary patterns of pathogens and their relationships with host species is critical to long-term efforts to control and mitigate their impact. Although a large body of research has emerged with a focus on understanding SARS-CoV-2 origins and the virus’ evolutionary trajectory (Andersen et al. 2020; T. Zhang, Wu, and Zhang 2020; Zhou et al. 2020), there has been less attention to host species, especially from a deeper evolutionary perspective. Given the importance of ACE2 in the pathogenicity of SARS-CoV-2 we focused this study on the molecular evolution of this protein in a wide range of mammalian species. Specifically, we wanted to test the Red Queen hypothesis, which postulates that an evolutionary arms race can force the pathogen and host to adapt new traits for their fitness and survival (Dawkins and Krebs 1979). We hypothesized that as a critical receptor of coronaviruses, ACE2 is involved in an evolutionary arms race that has led to evidence of positive selection in various positions within this protein across the evolutionary history of mammalian lineages.

## Methods

We retrieved 129 ACE2 coding genes from marsupial and placental mammals from NCBI’s Ortholog database (Table 1). Protein sequences were aligned using the COBALT alignment tool (Papadopoulos and Agarwala 2007), then mapped back onto the nucleotide sequences to produce a codon-based alignment. A reference tree was generated with the amino acid sequences in MEGA-X (Kumar et al. 2018) using simple neighbour joining (Saitou and Nei 1987). Distances were computed using a JTT matrix (Jones, Taylor, and Thornton 1992), allowing rate variation among sites with a gamma shape parameter of 1. The data were then analyzed using two different methods: adaptive Branch-Site Random Effects Likelihood (Smith et al. 2015) to find evidence for positive selection along different branches of the tree, as well as Mixed Effects Model of Evolution (Murrell et al. 2012) to detect episodic selection among codon positions. Both analyses were performed using HyPhy 2.5 (Kosakovsky Pond et al. 2020). We corroborated this analysis with known functional domains of the human ACE2 focusing on sites that interact with coronaviruses (see RefSeq NG_012575.1). Tree visualizations were created using iTOL (Letunic and Bork 2019)

**Table 1:**
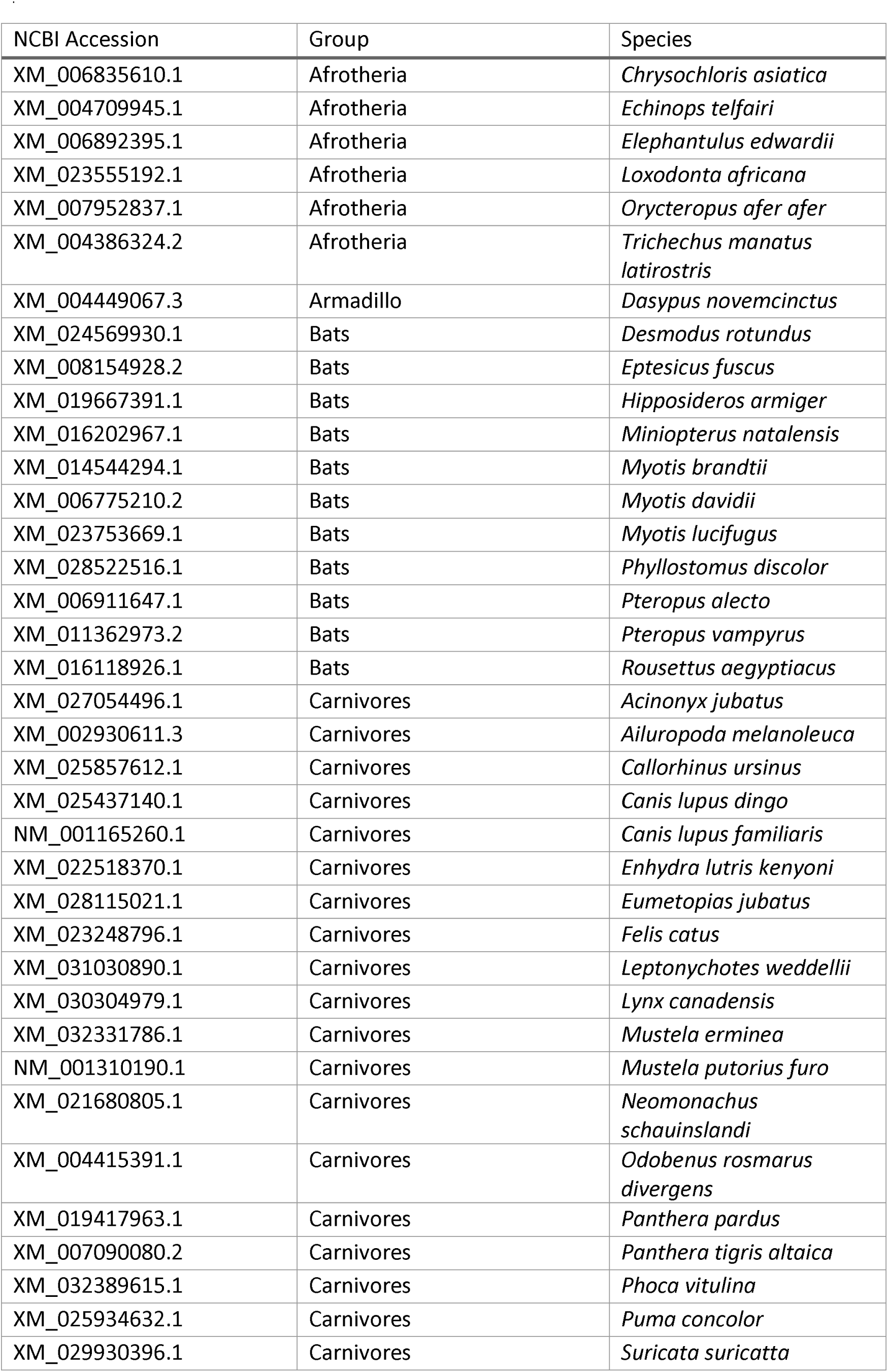
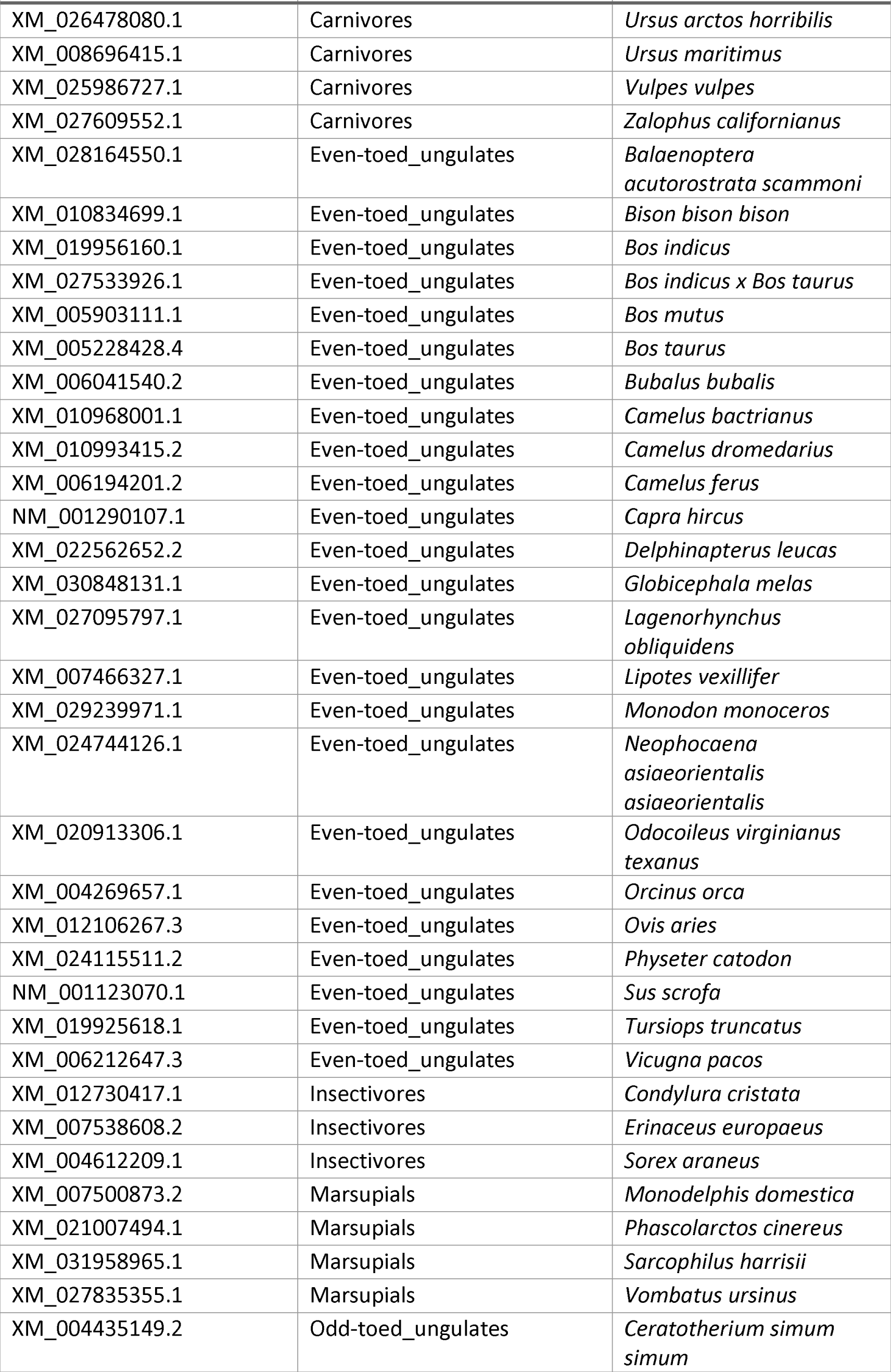
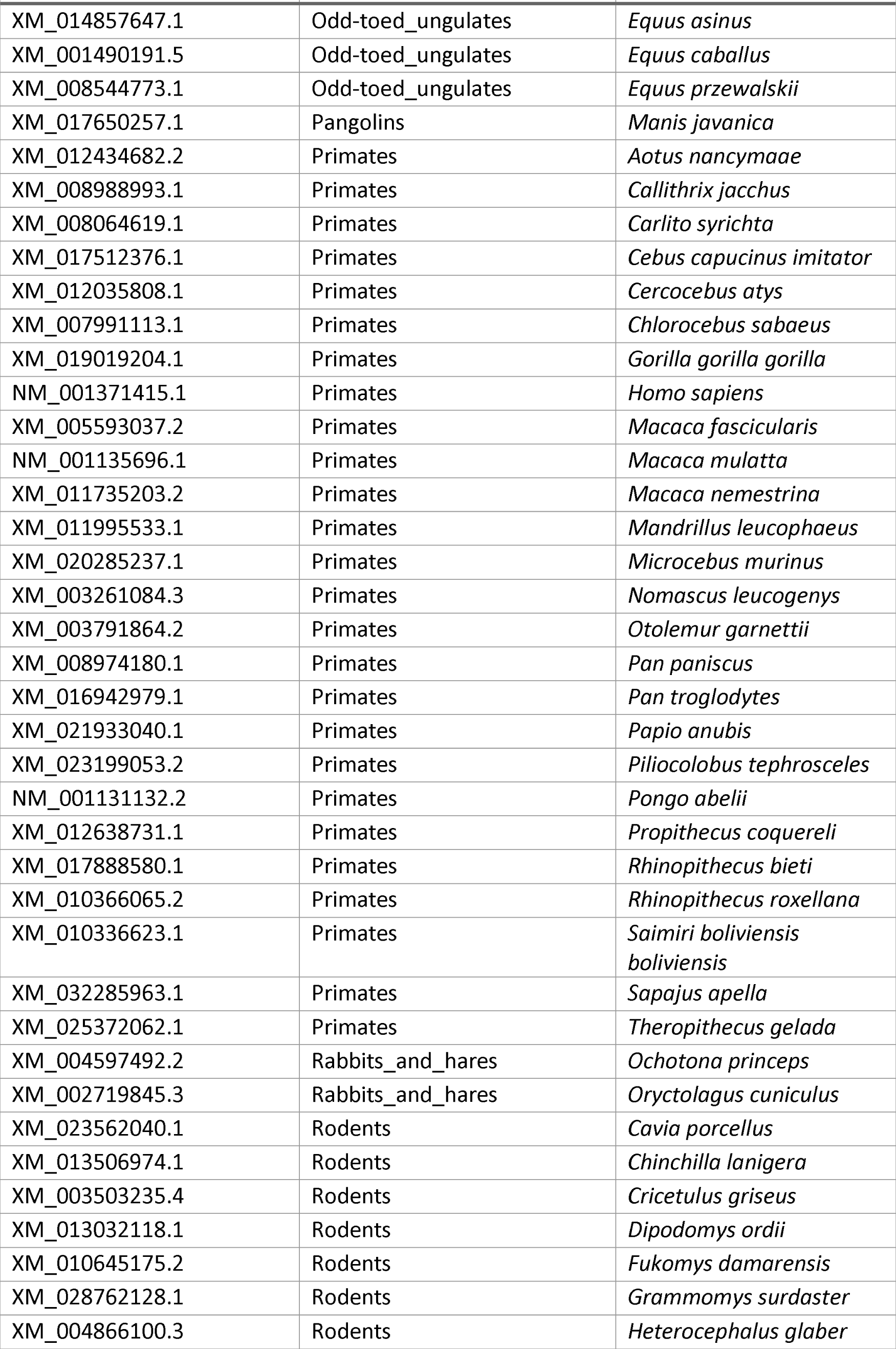
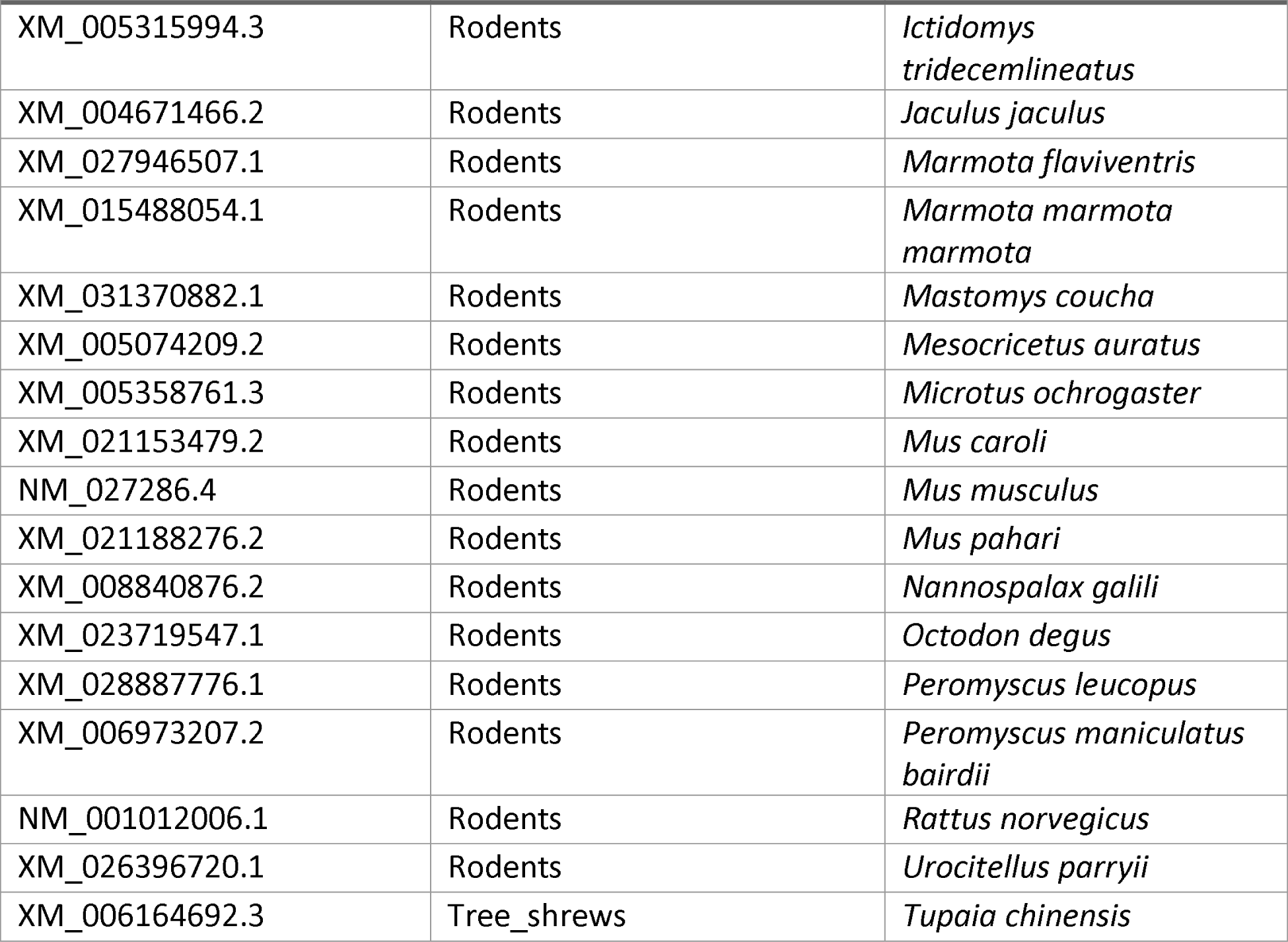
Sequences used in the analysis

## Results

Our analyses clearly show strong evidence of positive selection in ACE2 across the phylogeny of mammals at different phylogenetic depths. Figure 1 illustrates a phylogeny of ACE2 with cases that have undergone statistically significant positive selection (corrected p-value ≤ 0.05) shown in blue and red for moderate and strong positive selection, respectively (as judged by the proportion of sites with ω > 1). In some cases, deeper internal nodes in the phylogeny of ACE2 have undergone positive selection. For example, the basal branch of the primates shows evidence of positive selection. However, in other cases such as rodents or bats the evidence for positive selection is mostly found in shallower branches including a small number of species. And in some cases, such as cat and dog, the evidence is only at the species-level. A total of 13 species show positive selection at the species level and the strongest level of selection (i.e., greatest values of ω) is reflected in 5 of them. These include one marsupial species, the Tasmanian devil (*Sarcophilus harrisii*), donkey (*Equus asinus*), large flying fox (*Pteropus vampyrus*), Weddell seal (*Leptonychotes weddellii*), and dog (*Canis lupus familiaris*) (Figure 1). When considering the proportion of sites influenced by positive selection, these five species demonstrate different patterns with the largest proportion (6.6% of codons) in the seal and the smallest (0.4%) in the dog.

**Figure 1.**
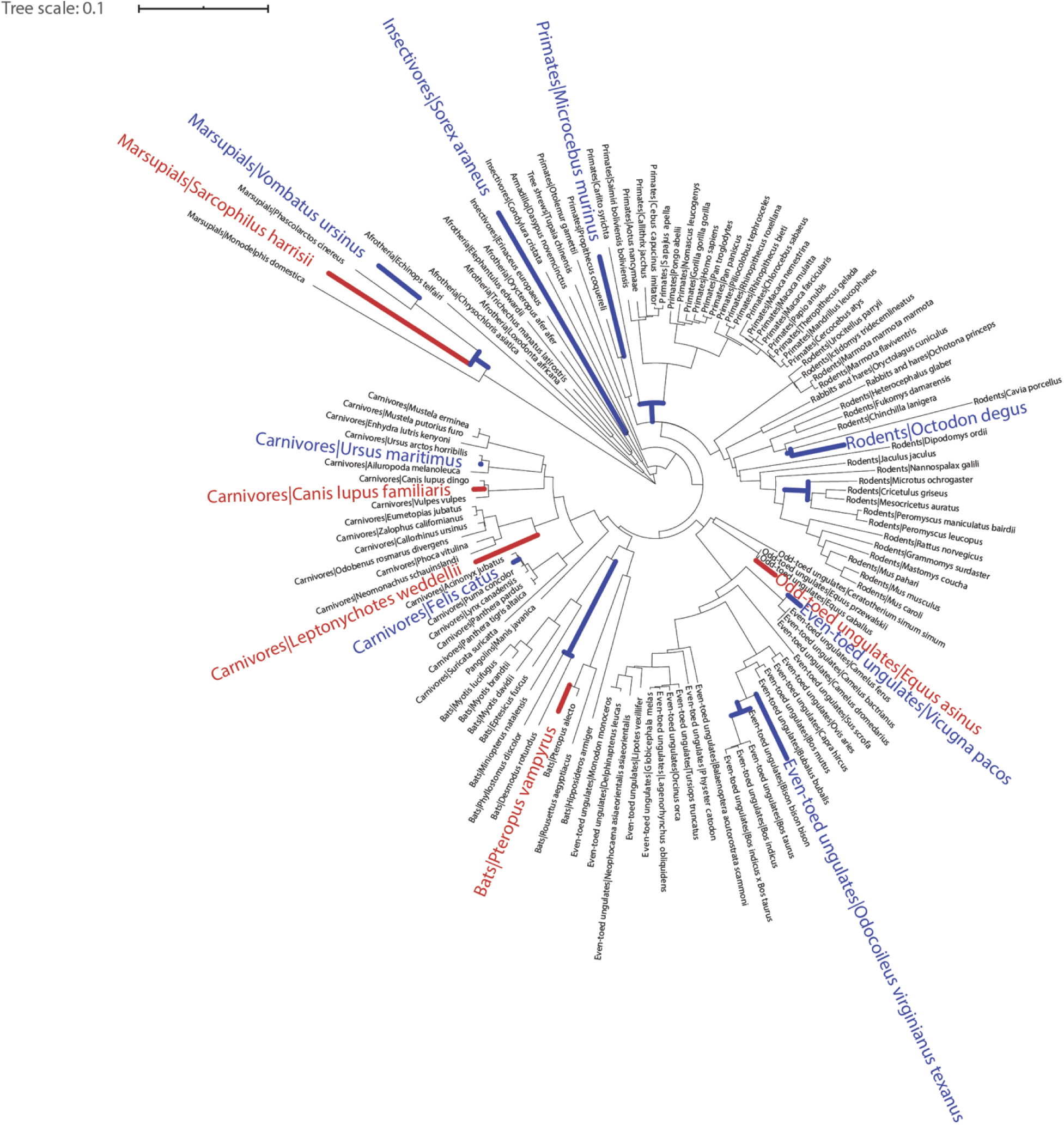
ACE2 protein in mammalian lineages is undergoing positive selection. Branches of the tree that are undergoing positive selection at various phylogenetic depths are color coded. Blue denotes moderate and red denotes strong positive selection.

In order to better understand the evolutionary trajectory of different amino acids in the ACE2 protein, we mapped the sites that showed episodic positive selection across the evolutionary tree to the ACE2 sequence in humans (shaded codons in Figure 2). There are 89 codons in all lineages that are under positive selection—approximately 10% of the sites in the protein. The phylogenetic distribution of these sites indicates up to 16 branches impacted by positive selection. When looking at the distribution of sites under positive selection, there are clear areas of concentration. The highest proportion of sites under selection belongs to a string of 30 codons starting at position 237, 14 of which (47%) show signs of positive selection among mammals.

**Figure 2.**
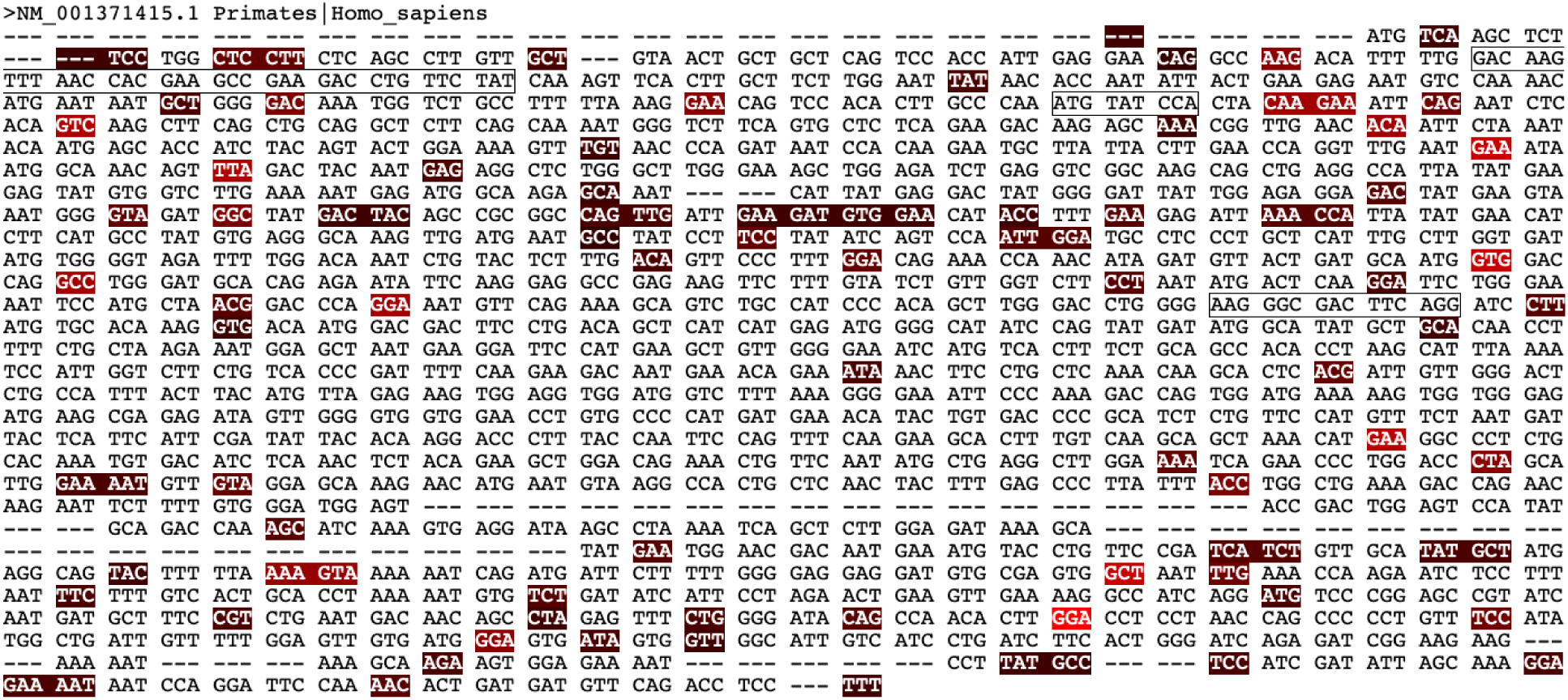
A substantial proportion of ACE2 codons are under positive selection in mammalian lineages. Nucleotide sequence of human ACE2 is used to map the position of codons under positive selection. Codons under positive selection are colored with darker codons show positive selection in more branches in the mammalian ACE2 phylogeny. The actual codons that are under positive selection are varied in different species tested. Three regions that are involved in binding with SARS-CoV-2 S1 protein are denoted in boxes.

An important consideration related to SARS-CoV-2 is the regions of the ACE2 protein where the viral spike S1 protein is known to bind, facilitating its entry into cells. Based on functional annotation from the human ACE2 protein, there are three regions in ACE2 that are known to interact with SARS-CoV and SARS-CoV-2 (boxed regions in Figure 2). While we do not see any codons under positive selection corresponding exactly to these protein regions in the human sequence, there are several codons that fall around these positions. We also note that there are several clusters of positive selection in ACE2 but no functional annotations for these sites are currently available.

## Discussion

Our results clearly show that ACE2 is undergoing positive selection in mammalian species and that a significant proportion of amino acid positions within this protein are affected. The primary function of ACE2 as a blood pressure regulator does not make it an obvious candidate for positive selection. However, ACE2 is the critical receptor for diverse and rapidly changing coronaviruses. This strongly supports the Red Queen hypothesis for an evolutionary arms race scenario where the ACE2—as a key trait of the host—is going through positive selection in order to escape/adapt to viral binding. To our knowledge, this is the first study that employs molecular evolutionary analyses to support an arms race that involves the ACE2 receptor in a wide range of mammalian species. Past research using structural biochemistry suggested that the Mouse Hepatitis Coronavirus (MHV) infection can lead to structural changes in its receptor by evolving a second allele indicating an evolutionary arms race (Peng et al. 2017). Here, we used a bioinformatics strategy from available sequences to demonstrate that the mammalian coronavirus receptor, ACE2, may be undergoing episodic evolution as a means of reducing the binding capacity of the viral spike proteins, lowering the probability or magnitude of infection.

As molecular evolutionary biologists, we provide a new perspective to a scientific body of work relevant to this global pandemic that has impacted nearly everyone on the planet. This is especially important since the current pandemic and previous SARS-CoV are linked to other mammalian species, chiefly bats (W. Li 2005; Y. Guan 2003; Andersen et al. 2020; Zhou et al. 2020). The conserved functional structure of ACE2 in mammals has made this protein an easy target for zoonotic viral pathogens. As demonstrated in our analysis, many lineages of mammals are evolving their ACE2, possibly in an attempt to escape the coronaviruses, which in turn can drive the evolution of new strains of these viruses. We can deduce that this phylogenetically-broad evolutionary arms race could potentially generate a very wide range of coronaviruses with heightened potential of jumping across species. As shown in several studies, close contact with species harboring these viruses can promote the transfer with devastating outcomes in recipient species such as the current pandemic.

